# Inferring plasticity rules from single-neuron spike trains using deep learning methods

**DOI:** 10.1101/2023.10.01.560360

**Authors:** Xingyun Wang, Jean-Claude Béïque, Richard Naud

**Affiliations:** Department of Cellular and Molecular Medicine, University of Ottawa, K1H 8M5, Ottawa, Canada; Centre for Neural Dynamics and Artificial Intelligence, University of Ottawa, K1H 8M5, Ottawa, Canada; Department of Physics, University of Ottawa, K1H 8M5, Ottawa, Canada

**Keywords:** plasticity rules, spiking neural network, deep learning

## Abstract

Synaptic plasticity is a core basis for learning and adaptation. Determining how synapses are altered by local signals – the learning rules – is the hinge about which brain activity pivots. A large number of in vitro characterizations have focused on restricted sets of core properties, but it remains to be established which if any of the known learning rules is most consistent with changes in activity patterns in behaving animals. To address this question, we hypothesize that the correlation between features of the activity of a single post-synaptic neuron and subsequent changes of the representations could be used to detect the underlying learning rule. Because this correlation is expected to be diluted in the notoriously large variability of brain activity, we test here learning rule inference based on passive observations of single neurons using deep artificial neural networks. Using simulated data, we found that both transformers, temporal convolutional networks, and SVM could classify learning rules far above the chance level, with transformers achieving the best overall accuracy. This performance can be achieved despite the presence of noise and representational drift. We further investigated the features used by the algorithms to perform the classification and found the deep net used inner temporal differences of distinct learning rules to separate learning trajectories. We also find, however, that the classification accuracy is sensitive to alterations in network properties. Our work illustrates that distinct learning rules’ generate distinguishable trajectories of responses, but warns against using simulation-trained classifiers to infer learning rules from real data.

## INTRODUCTION

Surviving and thriving are no easy tasks, but animals learn to adapt themselves to complex environments by reforming their neuronal networks. This ability is thought to arise from the interplay between learning rules, network architecture and the natural reverberation of neuronal activity implicated in adaptive behavior (1), where learning rules are a description of the main factors dictating the changes in synaptic strengths. Yet, despite their central role, we currently do not have a clear view of which learning rules are employed by different parts of the nervous system.

A number of learning rules have been put forward. The most common learning rules use the firing rate on both sides of the synapse to dictate plasticity. As such learning rules loosely encapsulate a postulate by Donald Hebb, these are referred to as Hebbian learning rules. Among such Hebbian learning rules, some formulations have been shown to learn selectivity to particular inputs (2–4), while others have been shown to emulate sophisticated forms of credit-assignment (5, 6). As the firing rate in these learning rules does not necessarily corresponds to the time-averaged rate of single neuron firing, but rather to a complex combination of activity in multiple neurons (7, 8) efforts to replace the firing rate by spike timings have been sought. In spike-timing-dependent plasticity (STDP), only one feature of the pre- and post-synaptic spike trains is influencing plasticity. The plasticity results from the relative coincidence between pre- and post-synaptic spikes (9–11). Another closely related example is burst-dependent plasticity (BDP), which focuses instead on the relative coincidence of different firing patterns such as single spikes and bursts or complex spikes in regulating plasticity (11–16). Recent work have exposed the potentially important alteration to BDP that occur when considering particularly large bursts, referred to as Behavioral Timescale Plasticity in cortex and hippocampus (BTSP (14, 17)). Beyond spike- and rate-dependence, some learning rules include neuromodulated-STDP (18–20) or oscillation-modulated plasticity (21). Going even more granular, molecular models of synapses are also types of learning rules. Here, no features of the pre- and post-synaptic spike train is extracted that is, neither rate or relative coincidence alone explains the plasticity. Instead, each spike triggers changes a cascade of electro-chemical events, which involve membrane potentials, intra-cellular calcium, activation of different kinases and other molecular cascades (22). Lastly, at the opposite degree of granularity, artificial neural network have learning rules that use detailed spatio-temporal information across the network. The backpropagation of error algorithm (BP) and its extension for temporal patternns, backrpopagation through time (BPTT) also hold different forms of learning rules.

Among this substantial number of learning rules, only some have been shown to be consistent with changes in representations as observed in vivo. In the hippocampus, a learning rule was inferred from in vivo observations during the emergence of place selectivity (23, 24). Two unique features of the hippocampus enable this discovery: a well-characterized structure of selectivity and a connectivity structure with weak recurrence. In the cortex, Lim et al. (25) revealed a signature of a learning rule by comparing the post-synaptic firing rate distribution between two learning sessions. This method is capable of unveiling a Hebbian-type learning rule during learning whose specific dependence on pre-and post-firing rate is inferred. Simulation using the inferred plasticity rule can reproduce experimental firing rate distributions of novel and familiar sessions. Importantly, Lim et al. could not discriminate which features of the pre-and post-synaptic spiking activity (spike time, bursts, etc.) were most consistent with the changes in firing rate distribution. Furthermore, this approach is insensitive to the trajectory of neuronal responses taking place between learned and naive states.

While many studies have established that some learning rules are consistent with a restricted set of features of neuronal activity (26–29), our work aims at finding which out of many is more consistent with a large amount of neuronal activity and thus with multiple features of the activ-ity. Asking a similar question entirely within artificial neural networks, Nayebi et al. (30) investigated machine learning methods that could successfully identify distinct optmization algorithms used to solve image classification tasks. In this work, the activation patterns predicted the learning rules with very high accuracy. This result is consistent with Portes et al. (31) which found a neural activity metric to distinguish supervised from reinforced learning rule in a recurrent neural network. Notably, Nayebi et al. (30) utilized a population of post-synaptic neurons to classify the learning rules while we focus on classification from single-neuron spiking activity.

To assess the validity of a learning rule using observational data, its manifestation during learning in the behaving animal must be distinguishable from other possibilities. In this paper, we ask whether artificial neural networks can be used to classify distinct learning rules from the evolving activity of recorded single neuron. We built a biological spiking neural network (SNN) and let the spike train evolve according to a number of learning rules: those found in the slice physiology literature (STDP and BDP) but also those used in the machine learning field (i.e., backpropagation through time, BPTT). The spike trains during training were concatenated to generate learning trajectories.

Two distinct scenarios of operations were considered, one where learning rules were set to increase the adequation between the end-representation and its power to solve MNIST. The other was where the end-representaiton was fixed across the different learning rules. We also asked, in the TCNN, which features of the spike train were used to perform the classification. While we could classify with reasonable accuracy the different learning rules, small changes to the simulation parameters (e.g. changes in the baseline firing properties) incurred large changes in the prediction accuracy. Together, our work indicates that while machine learning can be paired with simulation data to classify learning rules in real data, such inferences are highly dependent on the parameters used for simulation and on the learning rules used as a comparison.

## RESULTS

To estimate the feasibility of separating plasticity rules from observational spiking activity, we simulated a spiking neural network (SNN) with distinct synaptic plasticity rules and trained an artificial neural network to classify the types of plasticity rules. We focused on a restricted yet representative set of learning rules. Namely, we considered two spike-based biological learning rules (STDP, BDP) and two variants of a machine learning rule called spike-based BPTT. The two variants of the back-propagation through time (BPTT) algorithms differed in their cost functions: one using *L*1 loss (BPTT1) and the other using *L*2 loss (BPTT2). A fifth condition was added as a negative control (NC). We describe each learning rule in turn:

- **Spike-timing-dependent plasticity (STDP)** For STDP, the synaptic weight is determined by the relative spike co-occurrence between pre- and post-synaptic neurons. Our implementation follows Ref. (32) which models experimental observations at GABAergic and glutamatergic synapses (33–35), but is distinct from the common asymmetric forms of STDP (7). The synaptic weight is updated at every post-synaptic spike time, *t*_*post*_, according to the learning rule, Δ*W* (*t*_*post*_) = *η*(*x*_*pre*_(*t*_*post*_) −*x*_*th*_)(*W*_*max*_− *W*), where *x*_*pre*_ is an eligibility trace of pre-synaptic spikes (*S*_*pre*_) with a timescale set by *α*^−1^ and dynamics following *dx*_*pre*_*/dt* = −*αx*_*pre*_ + *S*_*pre*_. The parameter *x*_*th*_ is an eligibility trace threshold for eliciting long-term potentiation (LTP). The synaptic weights *W* are modeled as saturating at *W*_*max*_.
- **Burst-dependent plasticity (BDP)** For BDP, the neuronal response is assumed to be made of singlets or bursts with different effects on the plasticity. Bursts are spikes preceded by maximum one interval less than 16 ms, while singlets are the spikes preceded by an interval more than 16 ms. For each post-synaptic neuron, we form a burst train, *B*(*t*), and an event train, *E*(*t*). The synaptic weight change follows 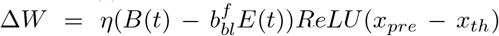*ReLU* (*x*_*pre*_ − *x*_*th*_) (11, 17, 36, 37), where 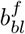 is a parameter regulating the strength of event-evoked long-term depression versus the strength of burst-evoked LTP. By controlling the burst fraction of the neurons, it is possible to control the plasticity (38). Thus, when setting the burst fraction *b*^*f*^ to the gradient of a loss, BDP can be used in a supervised setting. In what follows, we will use BDP to learn a particular representation **r**_*tar*_ using *Loss*_*BDP*_ = (**r**_*tar*_ −**r**_*BDP*_), where **r**_*BDP*_ is the representation observed in the simulations. This loss function determines how the neuron’s burst fraction changes, an action that is conceived as being mediated by inputs to apical dendrites (15, 38–41).
- **Spike-based surrogate gradient backpropagation through time (BPTT)** We used an artificial learning rule to train the network to achieve a target representation **r**_*tar*_. BPTT was used with surrogate gradients to pass the gradient through the nondifferentiable spike generation step (42, 43). We used two variants in order to test discrimination of learning rule parameters. For BPTT1, the SNN the loss function followed the *L*1-norm. For BPTT2, it is was instead an *L*2-loss. All other simulation parameters were identical between BPTT1 and BPTT2.
- **Weight updating fluctuations and representational drifts** In biological conditions, the synaptic weight updates are noisy, which might contribute to representation drift commonly observed (44, 45). For comparison, all plasticity rules (both biological and non-biological rules) have a noisy weight-updating process. Explicitly, the deterministic weight update Δ*W* is perturbed with a Gaussian noise, *ξ*_*W*_, having standard deviation *σ*_*W*_ .
- **Negative control (NC)** For the negative control, them weight update of the SNN is only driven by weight fluctuations, 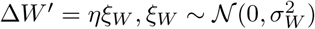.

These learning rules were implemented in SNN model that followed the architecture used by Diehl and Cook (32), as shown in Figure 1A (see Methods). In this network, the input layer sends Poisson spike trains representing images.

**Figure 1.**
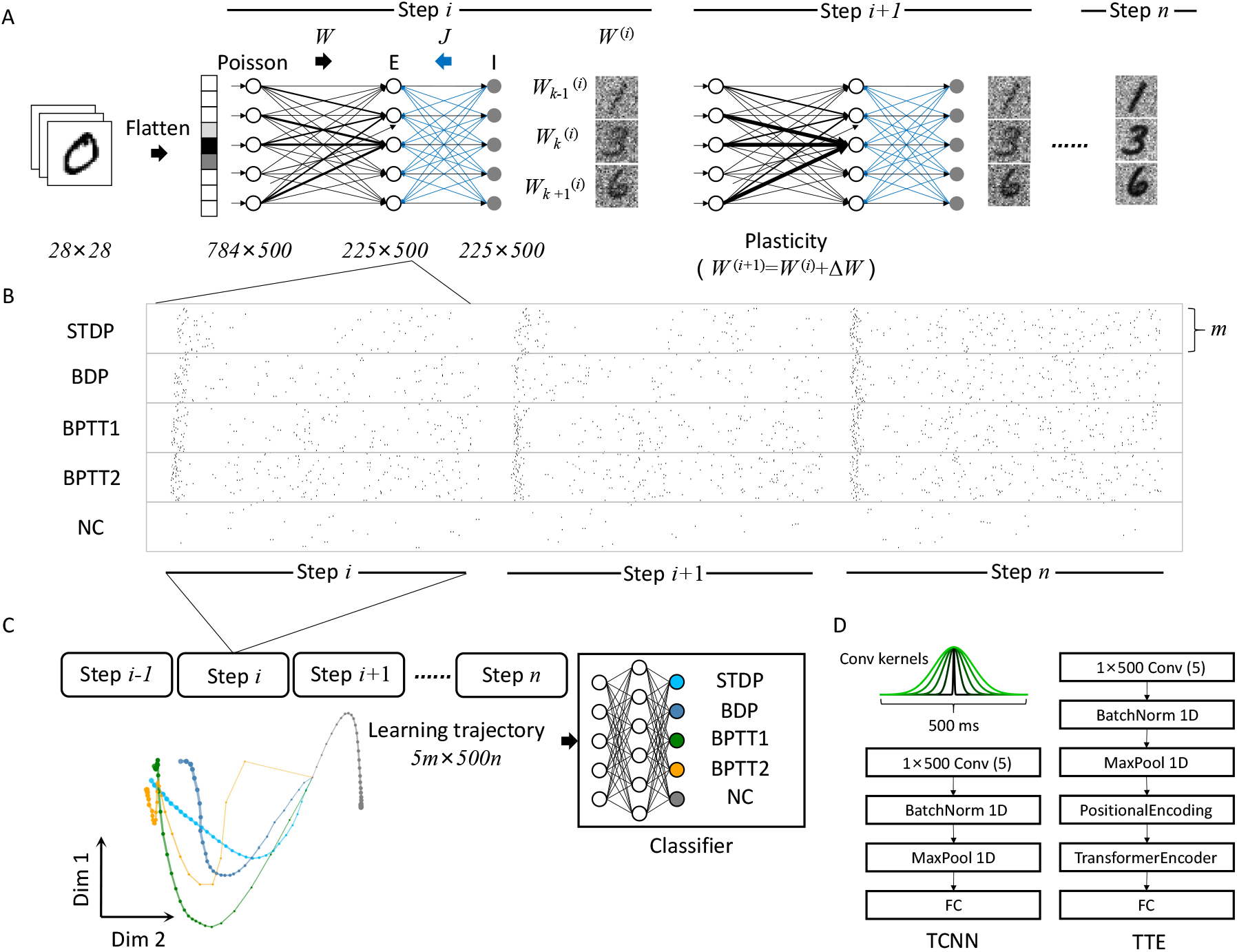
Trajectory generation from spiking neural network simulations in the context of image classification. **A** Schematic representation of the three layers of the SNN. Weights onto three exemplar neurons are shown as heatmaps, illustrating changes in weights between two learning steps.**B** During the learning process, for each of the 5 learning rules, *m* spike trains from a subset of responsive excitatory neurons are recorded for 500 ms and concatenated across *n* plasticity steps.**C** Concatenation of all learning steps of all responsive neurons form all the trajectories of the training set. Single spike train-learning rule pairs are fed into the classifier. Two-dimensional latent learning trajectories of distinct learning rules are also shown for visualization (using Gaussian process factor analysis, GPFA, see Methods). **D** Architectures of the two self-defined classifiers used. ‘Conv’ stands for convolutional and ‘FC’ stands for fully connected layer.

These spikes are sent to a layer made of excitatory cells with plastic weights, **W**, using one of the learning rules. Each excitatory neuron recruited one neuron providing lateral inhibition whose weights, **J**, were fixed. We chose this architecture because it has been shown to learn representations amenable to some degree of image classification with STDP. BDP (38, 46), BPTT1 and BPTT2 (47) can also be implemented in this architecture. Thus, choosing this architecture allowed us to compare the different learning rules within a similar context.

For each learning rule, we built trajectories of recorded responses to be fed into a classifier. To this end, we recorded the responses of all neurons in the network upon presentation of a restricted set of 300 examples from the MNIST dataset (48). While an example is being presented, the weights onto the excitatory neurons are updated according to one of the plasticity rules (Fig. 2A), such that the next presentation of the set of 300 training digits elicits slightly different responses. We repeated input presentation and plasticity for *n* plasticity steps. In the last plasticity step, we selected the subset of neurons having shown significant responses (firing rate above 6 Hz) for at least one of the ten categories of images in the dataset. Each responsive cell was thus associated with one or more category of image. We then extracted and concatenated the responses to that category for all plasticity steps (Fig. 2B), the result of which we call a trajectory. If a responsive neuron was responding to more than one category, then this neuron provided more than one trajectory. In this way, a single trajectory sees only one type of input category, the different trajectories contain information about all well-learned categories. As the trajectories contain information about the learning rules, the task of the classifier is then to map each trajectory onto the learning rule category (Fig. 2C). The classifier only sees the response of one neuron to one particular input category at a time. Different neurons or responses to different digits are treated as different examples.

**Figure 2.**
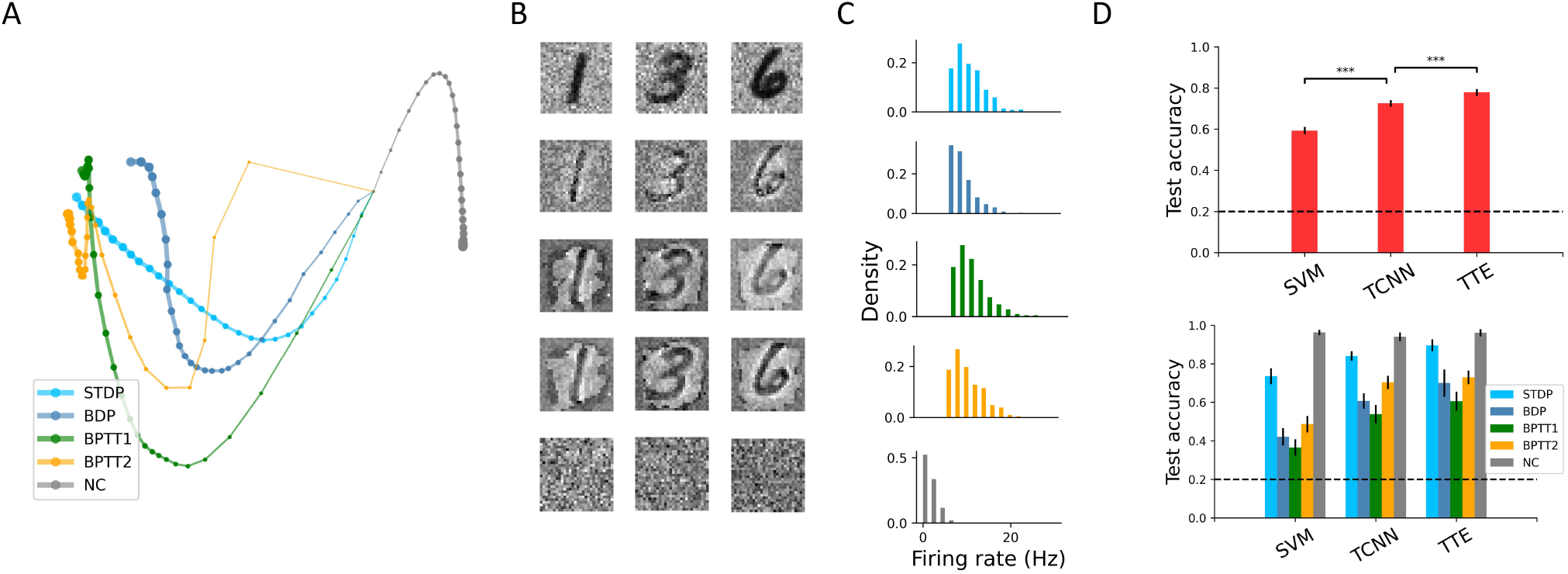
Learning rule categorization with fixed end-representation. **A** Latent space (two GPFA components) trajectories of distinct plasticity rules in the fixed end-representation protocol. The marker size of each learning trajectory grows with learning steps such that point of convergence with larger symbols on the left corresponds to the end representation.**B** Grayscale representation of weight values for connections onto three exemplar excitatory neurons (column-wise). From top to bottom, each row corresponds to a different plasticity rule: STDP, BDP, BPTT1, BPTT2 and NC. **C** Mean firing rate distribution accross responsive neurons (bin size: 2 Hz), with rows corresponding to rows in B.**D** Top, averaged overall accuracy based on 50 independent for each of the classifiers. Unpaired two-sample t-test, ^∗∗∗^*P* < 0.0001 is indicated. Chance level is indicated with a dashed line in each graph. Bottom, individual accuracy for each learning rules (different colors) according to the three classifiers considered.

### Single Neuron Responses can Reveal Learning Rules

First, we investigated the feasibility of separating trajectories while enforcing that the representation at the end of training was the same for the different learning rules, a protocol labeled ‘*fixed end-representation*’. This was introduced to disambiguate the role of the end-representation from that of the intermediate dynamics, we will illustrate the role of this constraint later in this article. Because all learning rules but STDP can be built to learn a particular representation, we have emulated the end-representation obtained under STDP plasticity using the other learning rules, setting **r**_*tar*_ = **r**_*ST DP*_, where **r**_*ST DP*_ is the response pattern under STDP after at the last plasticity step. In this scenario, the goal of BDP and BPTT learning rules is not, as is often the case, to accurately represent the different input categories.

Figure 2A shows a low dimensional projection of all trajectories, describing the evolution of activity patterns under different learning rules. For each learning rule, but not for our control condition, these network-level trajectories all end to one another. This is a natural consequence of the fact that we have constrained the end-representation to be the same. The changes in representation can be visualized further by inspecting the connection weights between the input neurons and individual excitatory neurons in the second layer (weights onto exemplar neurons are shown in Fig 2B). At the end of learning, STDP has developed a weight structure where shapes of particular digits can be visually recognized. Inspecting the end connection weights of corresponding neurons under BDP, BPTT1, and BPTT2 shows a similar weight structure. Similarly, for each learning rule, we can build the distribution of firing rates values observed across the network (Fig 2C). These are again very similar accross conditions. While the differences in learning rules is barely perceptible in the weights or in the firing rate distributions, the dynamics appear to reveal the different learning rules (Fig 2A).

We considered two classifiers to infer the plasticity types from the SNN trajectories:

- **Temporal convolutional neural network (TCNN)** (49) This classifier contains only one convolutional layer and one fully connected layer (Fig. 1D). The “temporal convolutional” means the TCNN convolves its one-dimensional kernels with each learning trajectory to extract temporal features. To improve the interpretability of the TCNN, we chose the shape of these kernels to be Gaussian with widths evenly tiled between 2 ms and 66 ms, as shown in Figure 1D left panel. These temporal kernels are fixed during training. We also made the bias in the convolutional layer and the fully connected layer 0, which lets the weight in the fully connected layer represent the contribution of each temporal feature for some plasticity category. During the classifier training procedure, only the weights in the fully connected layer are adjusted.
- **Temporal transformer encoder (TTE)**(50–52) The learning trajectory is first temporally embedded using a temporal convolutional layer, as in the TCNN. As a more complex classifier, information is then processed by positional encoding followed by a transformer encoder, which is cascaded by a fully connected layer (Fig. 1D).
- **Support Vector Machine (SVM)** As a comparison, we also implement SVM which is the benchmark of linear classifiers. All steps’ labeled learning trajectories from distinct learning rules are fed into Linear Support Vector Classifier (LinearSVC) model for training and inference.

After training our classifiers, we tested their accuracy on held out trajectories, with 50-fold repeats under independent initializations and training-test division. We used two accuracy metrics to measure classifiers’ performance: the first one is the overall accuracy, which is defined as the number of correct classifications divided by total number of samples; the second one is the individual accuracy, which is correct number of classifications for a given category divided by the sample number of this category, a quantity known as recall. We found that the overall accuracy is clearly larger than the chance level (0.2) for both classifiers (Fig. 2D). Using TTE, it reached 0.779, which is significantly higher than the that of TCNN (0.725; independent two-sample t-test, *P* < 0.0001). These results demonstrate that the learning rules can be inferred quite reliably from the response changes of individual neurons in the course of plasticity.

In the right panel of Figure 2D, we reported the individual accuracy for each plasticity rule (see also Table 1). We found that the overall accuracy is pulled down by BPTT1 and BPTT2, reflecting that distinguishing cost function is more difficult than distinguishing BDP from STDP. Nevertheless, learning trajectories from the same learning rules with distinct loss functions can be separated to some degree as the individual accuracy remains larger than chance level. We also found that classification is easiest for the STDP and NC category (see also the confusion matrices Fig. S2).. The classification patterns are roughly similar between classifiers, but for exception like BDP, which is best classified with TTE.

**Table 1.**
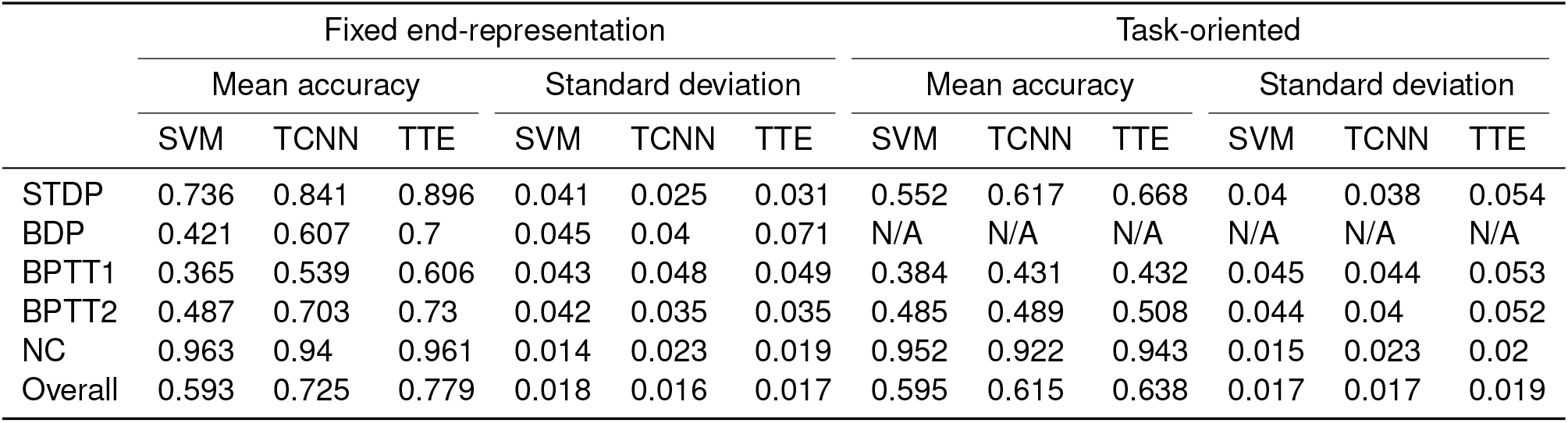
Prediction accuracy for different learning protocols.

|  | Fixed end-representation |  |  |  |  |  | Task-oriented |  |  |  |  |  |
| --- | --- | --- | --- | --- | --- | --- | --- | --- | --- | --- | --- | --- |
|  | Mean accuracy |  |  | Standard deviation |  |  | Mean accuracy |  |  | Standard deviation |  |  |
|  | SVM | TCNN | TTE | SVM | TCNN | TTE | SVM | TCNN | TTE | SVM | TCNN | TTE |
| STDP | 0.736 | 0.841 | 0.896 | 0.041 | 0.025 | 0.031 | 0.552 | 0.617 | 0.668 | 0.04 | 0.038 | 0.054 |
| BDP | 0.421 | 0.607 | 0.7 | 0.045 | 0.04 | 0.071 | N/A | N/A | N/A | N/A | N/A | N/A |
| BPTT1 | 0.365 | 0.539 | 0.606 | 0.043 | 0.048 | 0.049 | 0.384 | 0.431 | 0.432 | 0.045 | 0.044 | 0.053 |
| BPTT2 | 0.487 | 0.703 | 0.73 | 0.042 | 0.035 | 0.035 | 0.485 | 0.489 | 0.508 | 0.044 | 0.04 | 0.052 |
| NC | 0.963 | 0.94 | 0.961 | 0.014 | 0.023 | 0.019 | 0.952 | 0.922 | 0.943 | 0.015 | 0.023 | 0.02 |
| Overall | 0.593 | 0.725 | 0.779 | 0.018 | 0.016 | 0.017 | 0.595 | 0.615 | 0.638 | 0.017 | 0.017 | 0.019 |

**Table 2.** Perturbation results of different learning protocols.

|  | Fixed end-representation |  |  |  |  |  | Task-oriented |  |  |  |  |  |
| --- | --- | --- | --- | --- | --- | --- | --- | --- | --- | --- | --- | --- |
|  | Mean accuracy |  |  | Standard deviation |  |  | Mean accuracy |  |  | Standard deviation |  |  |
|  | SVM | TCNN | TTE | SVM | TCNN | TTE | SVM | TCNN | TTE | SVM | TCNN | TTE |
| STDP | 0.741 | 0.796 | 0.788 | 0.032 | 0.03 | 0.071 | 0.394 | 0.535 | 0.483 | 0.04 | 0.042 | 0.081 |
| BDP | 0.421 | 0.53 | 0.677 | 0.036 | 0.05 | 0.104 | N/A | N/A | N/A | N/A | N/A | N/A |
| BPTT1 | 0.335 | 0.386 | 0.385 | 0.043 | 0.044 | 0.09 | 0.271 | 0.446 | 0.466 | 0.046 | 0.051 | 0.073 |
| BPTT2 | 0.489 | 0.694 | 0.694 | 0.042 | 0.041 | 0.071 | 0.306 | 0.414 | 0.41 | 0.043 | 0.048 | 0.056 |
| NC | 0.951 | 0.935 | 0.969 | 0.016 | 0.018 | 0.016 | 0.968 | 0.885 | 0.943 | 0.013 | 0.027 | 0.019 |
| Overall | 0.589 | 0.669 | 0.702 | 0.018 | 0.015 | 0.023 | 0.483 | 0.569 | 0.575 | 0.022 | 0.023 | 0.025 |

### Inferring learning rules in a functional context

We next investigated a condition where the end goal of plasticity was not constrained to reach the same end representation. We defined this condition as the task-oriented protocol. As BDP and BPTT rules require a loss function, we made adaptations specific to each learning rule (see Methods). And, we didn’t consider BDP in this condition because it was hard to implement in a functional context.

In this scenario, we found that the classifiers could again successfully predict the learning rule categories (Fig S1D). The overall accuracy is higher than the chance level (0.25) for both classifiers (Table 1). The accuracy to individual learning rule indicates that it is led by reliable classification of the negative control. BPTT1 and BPTT2 remain the most difficult classes to disambiguate. This is further illustrated by the confusion matrices (Fig. S2).

### Interpreting features underlying the TCNN classifier

Based on the above results, learning trajectories from different plasticity rules are tolerably separable. An important question for using this methodology for answering neurobiology questions, however, is: what features of the neuronal responses are being used by the classifiers? We addressed this question with the TCNN because its simpler architecture makes this question tractable.

TCNN relies on temporal convolutions applied to the input spike trains. Each of the five kernels for these convolutions is shown in Figure 3. In these kernels, the narrowest (2 ms kernel width, Fig 3A; black curve) kernel filters for individual spike times. The slightly larger kernels can be sensitive to bursts of spikes over 10-40 ms (Fig 3A; green and dark blue curves). The largest kernels communicate a quantity consistent with a time-averaged firing rate. Since we fixed the convolutional layer (kernels and biases) and set fully connected layers’ biases to zero during learning, the weight of the fully connected layer indicates the relative contribution of each temporal kernel to each digit category. Investigating what parameters the learning algorithm has chosen can reveal what feature is relevant to the classification. Note that there is a set of weights for every time bin in the trajectory, so the relative relevance of each feature (or kernel) can change in the course of the learning trajectory.

**Figure 3.**
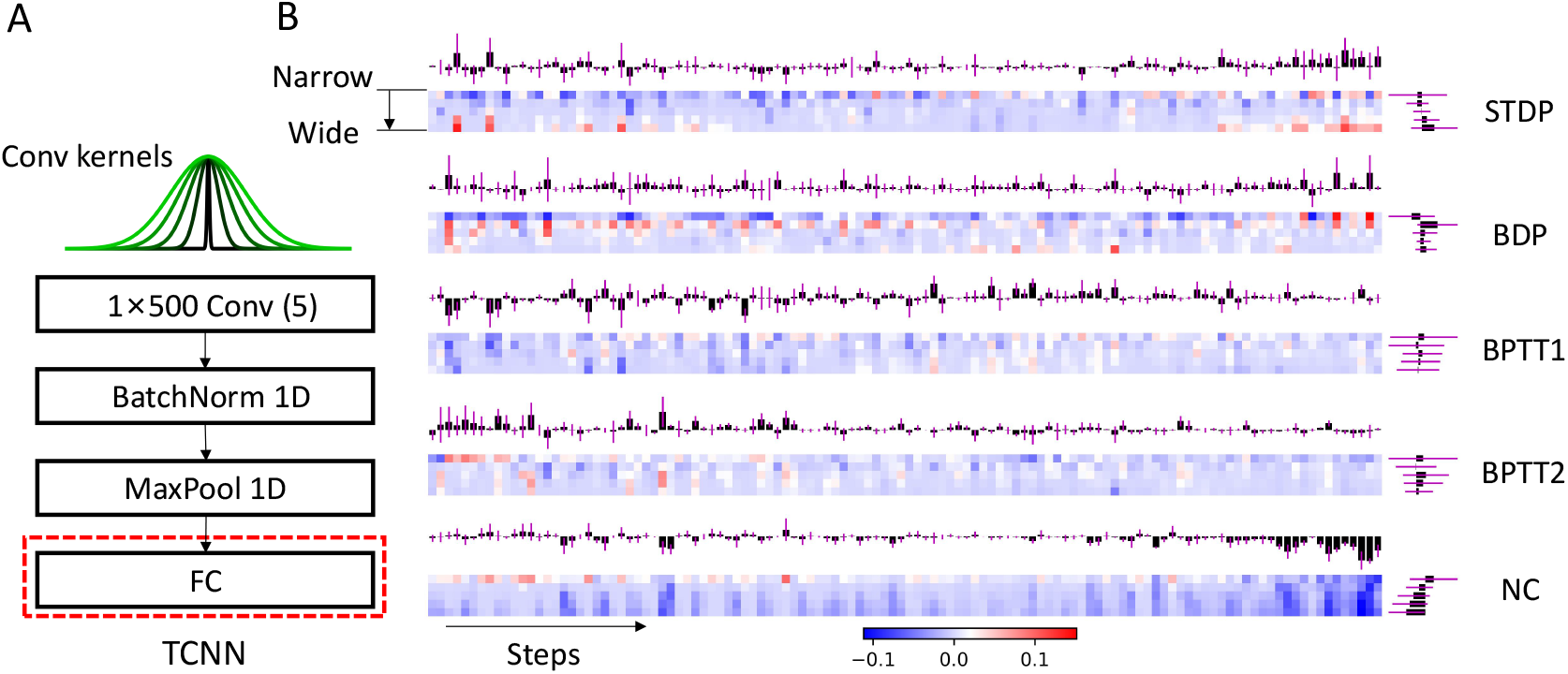
TCNN relies on different temporal features to predict learning rule identities. **A** Five Gaussian temporal convolution kernels are used in the TCNN. The architecture of the TCNN is shown below the kernels, whereby the convolutional slayer feeds in a batch-norm operation, then a maxpool operation followed by a fully connected (FC) connectivity to the readout layer. **B** Heat map of the fully connected layer’s weights (average value across 50 initializations) from each channel (lines) and each time unit (columns) for each learning rule. The column-wise (above the heatmaps) and line-wise (right of the heatmaps) weight average and standard deviation are shown.

Figure 3B displays the learned weights over the trajectory for each kernel width and each learning rule. To gain insight into the relevance of a temporal feature for classification, we computed the average weight for each kernel width. We found that NC is reported with a net positive contribution only for the most temporally precise kernels. NC is also distinguished by relatively low weights for the widest kernel towards the end of learning steps (Fig 3B; more negative weights on the right side). BPTT1 and BPTT2 rely on the positive contribution of the most temporally precise kernels of specific timings. Consistent with the larger proportion of bursts in BDP, this learning rule is distinguished with an intermediate kernel width and a negative contribution from a temporally precise kernel. The STDP relies on a negative contribution from the temporally precise kernel at the beginning and positive contributions from the rate-computing kernels in the end. These observations help provide an explanation of how the classifier is able to discriminate plasticity rules.

### Dependence on Simulation Parameters

Having established that plasticity rules can be identified based on how a single neuron changes its responses through learning, we then asked if our such trained classifiers could in principle be applied to recordings of single neurons through learning. We thus conducted ‘out-of-distribution tests’ whereby the classifiers were tested using simulation parameters distinct from those used in training. We considered two types of perturbations: 1) Burst fraction perturbation, where we trained with sparse burst fraction baseline (*b*_*f*_ = 0.2) and tested with a higher one (*b*_*f*_ = 0.4; Fig 4A). 2) Background perturbation, where we set the noise level to 0.05 in training and to 0.1 in the test set (Fig 4B). The first perturbation was applied to the fixed end-representation protocol, while the second perturbation was applied to the task-oriented protocol.

**Figure 4.**
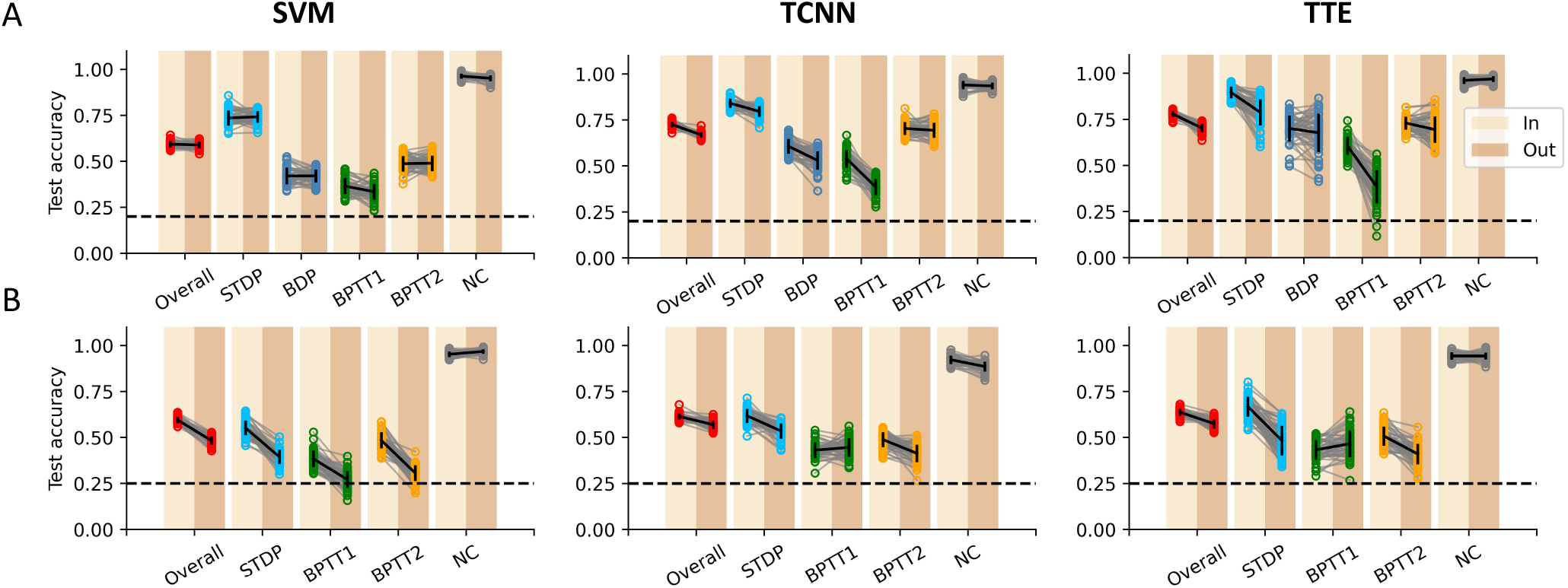
Plasticity prediction in out-of-distribution learning trajectories. **A** For the fixed end-representation protocol, the SVM (left panel), TCNN (middle panel) and TTE (right panel) are tested with neurons either having the same burst fraction baseline as in the training set (left points in a pair) or increasing the burst fraction baseline to 0.4 (right points in a pair). **B** Same as A but for background noise perturbation in the task-oriented protocol.

As shown in Figure 4, the total prediction accuracy under the fixed end-representation protocol decreases under burst fraction perturbation. For all classifiers, the overall mean accuracy is affected. This decrease in accuracy arises in large part from the classification of the BPTT1 learning rule. Perturbation of the background firing rate also caused a significant overall accuracy decrease for all classifiers. Again this serious decrease in accuracy arises in large part from the classification of the BPTT1 and BPTT2 learning rules, but present for all learning rules. Although these detrimental effects could be mitigated by training with a greater diversity of simulation parameters, the fact that simulations are bound to differ in terms of many parameters, the compound effect of such mismatches is likely catastrophic to this approach.

## CONCLUSION

To verify the feasibility of inferring biological plasticity rules by observing single-neuron spike trains, we trained multiple SNN models to classify images using distinct plasticity rules. The prediction accuracy for both learning protocols was largely above chance, despite the presence of representation drift and background noise. Different temporal components of the response are exploited by the machine learning algorithms to distinguish plasticity rules in a manner consistent with expectations. We found, however, that the methodology tended to be sensitive to the details of the simulation used to train the classifiers.

## METHODS

### Spiking neural network model

As described in the previous section, the SNN model has three distinct functional layers. The first layer is the pre-processing layer, which is designed to map the intensity of the image pixel into 500 ms lasting Possion spike train. The second and third layers are responsible for information processing. Excitatory neurons in the second layer generate the tuning for the digit features using the learned feedforward connection. And, inhibitory neurons in the third layer, which are point-wisely triggered by previous excitatory neurons, induce competition among excitatory neurons by the fixed lateral inhibition connection.

#### Possion spike train coding for digit stimuli

Each stimulus is a digit picture sampled from the MNIST dataset, in which every digit’s pixel intensity value is ∈ [0, 1]. It will first be flattened into a 784 × 1 vector and feeds to the first layer in a one-to-one manner. In the first layer, a single pixel’s intensity will be multiplied by a modified factor of 0.25 and then be used to generate the Poisson spike train. Firing rates of the generated spike train are from 0 to 288 Hz.

#### Single neuron model in the second and third layer

Both excitatory neurons in the second layer and inhibitory neurons in the third layer are described as the noisy Leaky Integrate-and-Fire (LIF) model, which has the classic form as

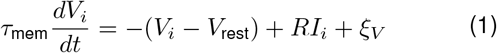

where *V*_*i*_ is the membrane potential of the neuron *i, V*_reset_ = 0 is the reset membrane potential, *I*_*i*_ is the input current for neuron *i, R* is the resistance for the input current, *τ*_mem_ is the time constant for the membrane voltage decay, *ξ*_*V*_ is the background Gaussian noise. The input current *I*_*i*_ can be modeled as the integration of the feedforward and the recurrent current. Mathematically, we describe it as

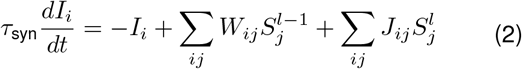

in which *W*_*ij*_ is the feedforward connection from neuron *j* to neuron *i*, 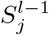 is the spike train of previous layer *l* – 1 neuron *j, V*_*ij*_ is the recurrent connection from neuron *j* in the same layer *l*, 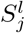 is neuron *j*’s spike train. For the LIF model, the spike train *s*_*i*_ is

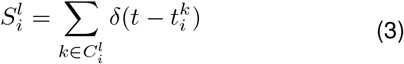

Here 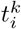 is the timing of *k*th spike for neuron *i*, and 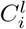 is the set of *l* layer neuron *i*’s spike timing. Typically, the spike is generated by a nonlinear Heaviside function Θ

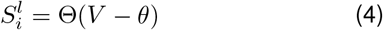

where *θ* is the spiking threshold and is set to be 1.0. To enable surrogate gradient calculation, we use a sigmoid function in the backward step to replace the Heaviside function

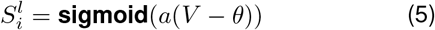

if *a* is large enough (i.e., *a* = 100), the output can be approximated as a Heaviside function and differentiable. For simplification, the excitatory neuron in the second layer is re-written as

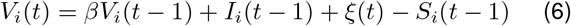

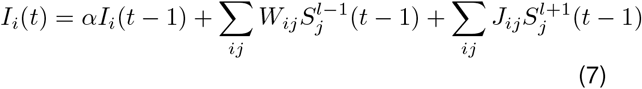

where 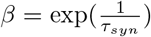 and 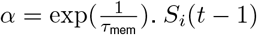 works for resetting the membrane potential after neuron firing. Here, the term 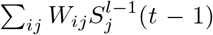 represents linear integration for the feedforward Poisson spike train. The term 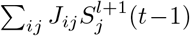 denotes the global lateral inhibition from the third layer instead of the recurrent connection within the second layer, and *J*_*ij*,*i*=*j*_ = 0. Considering the homeostasis, which limits the firing rate of the single neuron, we induce a drift threshold

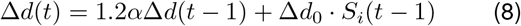

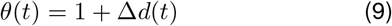

where Δ*d*_0_ is the initial drift of threshold after neuron firing, Δ*d*(*t*) will gradually decay to 0. Therefore, the threshold *θ* will decrease back to 1.0 in a while. Similar to the excitatory neurons, the membrane potential of inhibitory neurons in the third layer obeys the same rule as Eq. 6, but the input current *I*_*i*_ should be described as

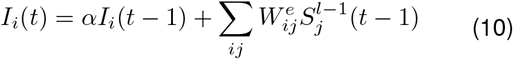

where 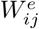 is point-wise poistive feedforward connection from the excitatory neurons, 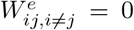. And, the drift threshold of the inhibitory neurons obey

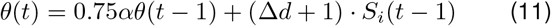

#### Parameters initialization

The learned feedforward connection to the second layer excitatory neurons is initialized by sampling from a uniform distribution *𝒰* (−1*e*^−3^, 1*e*^−3^). The point-wise connection value is set up as 1.0, and the lateral inhibition value for each excitatory neuron is assigned as −0.005. We determine *τ*_mem_ = 500ms, *τ*_syn_ = 5ms and Δ*d*_0_ = 0.2. The background noise *ξ*_*V*_ (*t*) ∼ 𝒩 (0, 0.05^2^).

#### Weight updating fluctuation and representational drifts

In biological conditions, the synaptic weight updates are noisy, which might contribute to representation drift commonly observed (44, 45). For comparison, all plasticity rules (both biological and non-biological rules) have a noisy weight-updating process. Here, we perturbed the deterministic weight update Δ*W* with a Gaussian noise, *ξ*_*W*_, having standard deviation *σ*_*W*_ such that: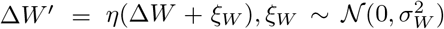, where *η* is the learning rate. During all the learning steps, we set the fluctuation level to be half of the maximum value of the initially learned feedforward connection strength, *σ*_*W*_ = 5*e*^−4^.

#### The task-oriented protocol

Here, SNNs trained with different plasticity rules must complete the MNIST task independently, with adaptations specific to each learning rule (except BDP which cannot be adapted to this protocol):

- **STDP** The SNN representation evolves without supervision. Previous work demonstrated that this change in representation is associated with improvement on MNIST when paired with a readout mechanism (32).
- **BPTT1 and BPTT2** We added a linear readout layer **W**_*lin*_ to map the excitatory neuron’s activity **r** into the output category. The readout layers for both BPTT1 and BPTT2 are fixed in order to constrain the weight updates to only occur in the weights from the input layer. This requirement prevents different loss functions from affecting the readout weights, thus enforcing a more direct effect of the loss function onto the observed trajectories. BPTT1 uses smooth *L*1 loss to the one-hot representation **y** of categories, which is *Loss*_*BP TT* 1_ = *smoothL*1(**W**_*lin*_**r, y**). BPTT2 uses classic *NLLLoss* loss to learn the mapping from input to target categories, which is written as *Loss*_*BP T T*_ = *NLLLoss*(*Softmax*(**W**_*lin*_**r**), **y**).

#### Gaussian Process Factor Analysis (GPFA)

To visualize learning trajectories from different learning rules, we used GPFA to show each learning rule’s learning trajectory in two dimensional space. We first used *n* spiking learning trajectories of each learning rule as instantaneous firing rate (with correct alignments) combined with inhomogeneous Poisson process to re-generate spike trains for 60 trials. After that, for each learning rule, we would get 60 × *n* Poisson spike trains. For each learning rule, we fed those spike trains into GPFA fitting function with 500 ms time bin to get each trial’s 2*D* trajectory of 30 steps. We showed mean trajectory of trials for each learning rule after subtracting the value of the starting step to make it begin from [0, 0] in 2*D* space.

#### Computing resources

This work was done by using Google Colab. All SNN simulations were run on the high-ram CPU runtime. Classifiers were trained on the NVIDIA Tesla T4 GPU.

## ACKNOWLEDGEMENTS

We wish to acknowledge that this work was carried out on the unceded and unsurrendered land of the Algonquin Anishinaabe people.

The authors are grateful for funding. This work was funded by CIHR Project Grant 175319, CIHR Project Grant 175325, as well as NSERC Discovery Grant RGPIN-2024-05162.

## AUTHOR CONTRIBUTIONS

XW and RN designed the study. XW performed the simulations and analyzed the data. RN and JCB provided funding. JCB provided criticism on the manuscript. RN and XW wrote the manuscript.

## Notes

https://drive.google.com/drive/folders/1-f2EmMORZeYs-h9u71eRiTzyQM-zkbPX?usp=drive_link

### Competing Interest Statement

The authors have declared no competing interest.

### Summary of Updates

We rewrite the paper into the journal paper format. The main result is not changed.

