## Supplementary material for "Inferring plasticity rules from single-neuron spike trains using deep learning methods"

### A SUPPLEMENTARY FIGURES

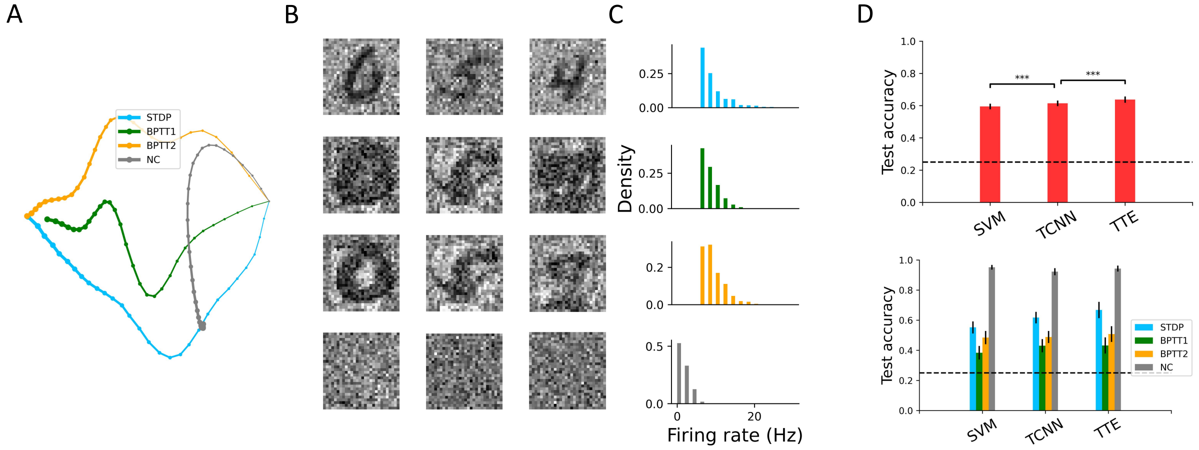


**Figure S1. Learning rule categorization in the task-oriented protocol.** As in Fig. 2 but the simulations are letting BPTT1 and BPTT2 learning rules attempt to classify the input categories.

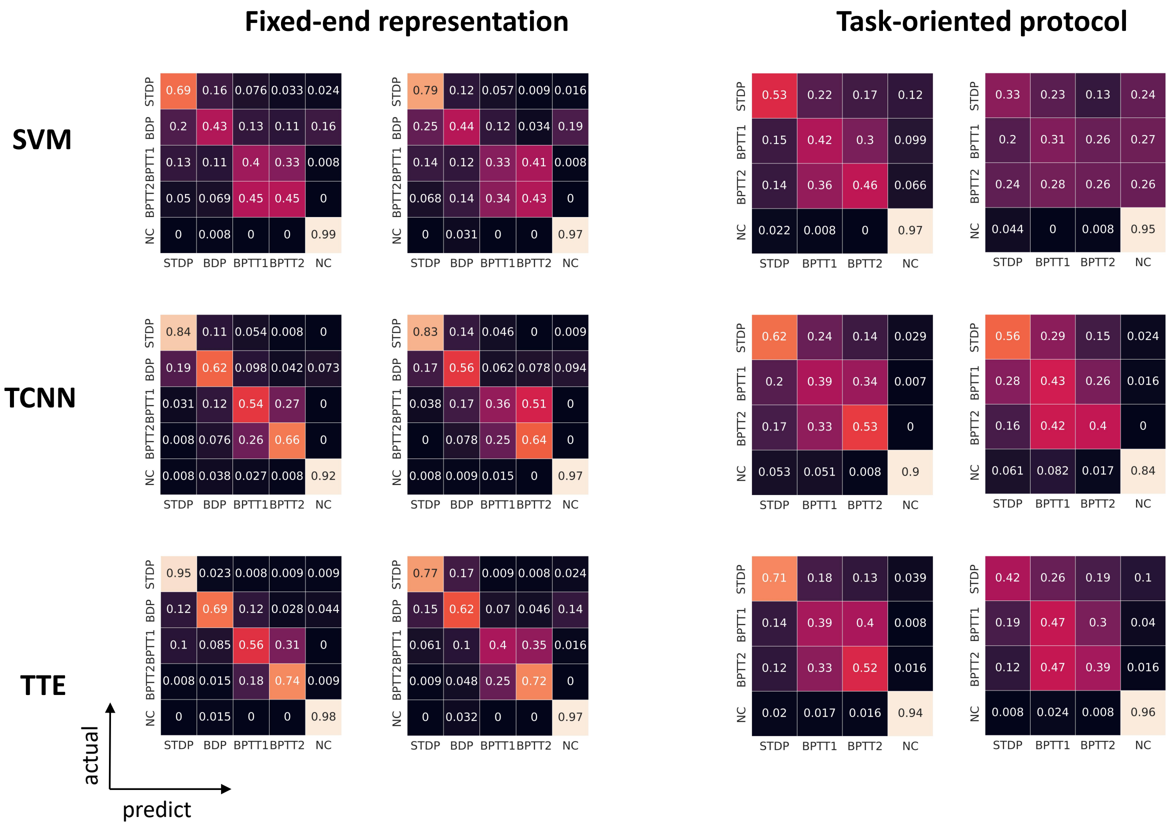


**Figure S2. Confusion matrix for in-distribution and out-distribution test.**
